# Pharmacological enrichment of polygenic risk for precision medicine in complex disorders

**DOI:** 10.1101/655001

**Authors:** William R. Reay, Joshua R. Atkins, Vaughan J. Carr, Melissa J. Green, Murray J. Cairns

## Abstract

Individuals with complex disorders typically have a heritable burden of common variation that can be expressed as a polygenic risk score (PRS). While PRS has some predictive utility, it lacks the molecular specificity to be directly informative for clinical interventions. We therefore sought to develop a framework to quantify an individual’s common variant enrichment in clinically actionable systems responsive to existing drugs. This was achieved with a metric designated the *pharmagenic enrichment score* (PES), which we demonstrate for individual SNP profiles in a cohort of cases with schizophrenia. A large proportion of these had elevated PES in one or more of eight clinically actionable gene-sets enriched with schizophrenia associated common variation. Notable candidates targeting these pathways included vitamins, insulin modulating agents, and protein kinase inhibitors with putative neuroprotective properties. Interestingly, elevated PES was also observed in individuals with otherwise low common variant burden. The biological saliency of PES profiles were observed directly through their impact on gene expression in a subset of the cohort with matched transcriptomic data, supporting our assertion that this framework can integrate an individual’s common variant risk to inform personalised interventions, including drug repositioning, for complex disorders such as schizophrenia.

## INTRODUCTION

A significant burden of disease is caused by complex traits including psychiatric and neurobehavioural disorders, inflammatory and autoimmune disorders, metabolic and cardiovascular disease, and cancer. Until relatively recently it was difficult to identify the heritable components of these traits, however, the emergence of well powered genome-wide association studies (GWAS) using large cohorts assembled by collaborative consortia are revealing important insights into their common variant architecture ^1^. While collectively this information has been vital to map genes and pathways that are likely to be etiological factors, the small effect size of each variant, and their heterogeneity in the population make their relevance to individuals with the disorder highly variable, relatively specific, and fairly minor with respect to the total variant burden. They also present as relatively small targets for therapeutic intervention and may not attract the investment needed for pharmaceutical development. We therefore need mechanisms for using this vast amount of diverse genetic information to maximise its utility for therapeutic advances. This requires a personalised approach that can capture variant burden in affected individuals with respect to biological components that align with existing medications, and/or pathways of relevance to key pathophysiological processes, to provide sufficient support for the development of new interventions.

While approaches that summate the genomic risk burden in individuals, such as polygenic risk scoring (PRS), have demonstrated some predictive utility for complex traits (such as neuropsychiatric disorders ^2; 3^, diabetes ^4; 5^, cardiovascular disease ^6; 7^, and inflammatory disorders ^8; 9^) their composition of heterogeneous risk factors lack the biological salience needed to design a precision treatment strategy. We, however, hypothesized that the biologically supervised enrichment of trait-associated common variants in clinically actionable pathways would provide a means of pharmacologically annotating PRS in individuals. To test this proposal, we devised a statistical framework for scoring polygenic risk (at the multivariable level) in pathways relevant as therapeutic targets for complex traits. This quantitative approach, designated the *pharmagenic enrichment score* (PES), was developed to provide an indication of an individual’s exposure to risk variants that are potentially treatable by existing pharmacological agents, including many that have never been considered or tested previously for the condition/disorder they are experiencing. By focusing on biological pathways with known drug targets, we endeavour to enhance the clinical utility of polygenic risk approaches by providing novel and specific opportunities to identify treatment targets and/or repurpose existing drugs. This application of genome-wide common variant genotyping should have particular relevance for the precision treatment of individuals that are resistant to currently indicated medications. In this study we outlined the PES approach and sought to exemplify its utility in individuals with the complex psychiatric condition, schizophrenia.

## MATERIALS AND METHODS

### Pipeline for the derivation of *pharmagenic enrichment scores*

The methodology developed for constructing *pharmagenic enrichment scores* (PES) is outlined in a schematic presented in Supplementary Figure 1. We exploit the results of gene-set association analysis to aggregate variants from GWAS into gene-sets which may be candidates for pharmacological intervention. These gene-sets were high quality canonical and hallmark pathways sourced from the molecular signatures database (MSigDB) ^10^. Pathways were designated as clinically relevant if they contained at least one gene annotated to interact with an approved pharmacological agent in the DrugCentral database as classified by the Target Central Resource Database (TCRD, genes annotated as T_clin_) ^11^.

Firstly, this process tests the combined effect of variants at the gene level. We utilised an omnibus *P* value test to achieve this, whereby a linear combination of variant-wise *P* values in a gene is compared to a null distribution to derive a combined *P* value for that gene. This was performed using the MAGMA package to account for linkage disequilibrium between variants in the approximation of the null *χ*^2^ distribution ^12^. We mapped SNPs to protein coding genes (hg19, NCBI) with the genic boundaries encompassing 5kb upstream and 1.5 kb downstream to capture variation in regulatory regions. Genes within the major histocompatibility complex (MHC) on chromosome 6 were excluded from this study due to the complexity of haplotypes in that region. Traditionally, the full breadth of variants available in the summary statistics are utilised in these models. This, however, may miss important aspects of the biological interpretation of the genomic signal. PRS derived from large GWAS cohorts are tested at different P value thresholds (*P*_T_) to exploit a model which explains the most variance between the case and control groups (in a dichotomous construct). For instance, a *P*_T_ < 0.05 means that only *P* values in the GWAS with a stronger association (*P* value) than this threshold are included. A range of *P*_T_ have been suggested to be optimal for PRS in several disorders depending on their genomic architecture ^2; 7; 13^. Aggregating the combined effects of variants in genes at different *P*_T_ may therefore capture the biological complexity of the signal at varying degrees of polygenicity.

Once variants are aggregated in genes at varying significance thresholds (*P*_T_), we conduct gene-set association of the pre-defined druggable gene-sets with the trait of interest. This was a competitive association test, which tests the null hypothesis of genes in the set being no more strongly associated than all other genes. This model was implemented with the MAGMA package. Enriched pathways with a known drug target uncovered from this pipeline then form the basis of calculating the *pharmagenic enrichment score* (PES). To profile individuals for a PES, genes which compromise the candidate pathway are extracted and cumulative genomic risk calculated in an analogous fashion to PRS, assuming an additive model ^2^. Each individual profiled thus has a genomic risk score within each clinically actionable pathway.

### Application of *pharmagenic enrichment score* to identify drug repurposing candidates for schizophrenia

To identify clinically actionable gene-sets and construct PES, we processed the 2014 psychiatric genomics consortium GWAS for the complex neuropsychiatric disorder schizophrenia with the pipeline described above ^3^. Variants were selected based on their significance (*P* value) for inclusion in the at four different thresholds – all SNPs, *P* < 0.5, *P* < 0.05, and *P* < 0.005. Geneset association was conducted on 1012 MSigDB pathways with at least one druggable (T_clin_) gene. To capture a wide variety of pathways for a complex phenotype like schizophrenia, we used a nominal significance threshold of *P* < 0.001 to select candidate pathways for PES.

We annotated each of these candidate pathways for their drug interactions, tissue specificity, and phenotypic associations. Using WebGestalt, drugs which target a statistically significant number of genes in the PES gene-sets were identified after the application of multiple testing correction (FDR < 0.05) ^14^. A minimum overlap of at least three targets overlapping the geneset for each pharmaceutical agent was also implemented. Drugs were also mapped to gene-sets using DGidb v3.02, with the top FDA approved drug per pathway was selected based on the DGidb score of interaction confidence between a T_clin_ gene and drug ^15^. Further, the tissue specificity of expression for all genes from the eight pathways was investigated using GENE2FUNC application of FUMA ^16^. Transcript expression in each of the 53 tissue types in the GTEx v7 dataset for the input pathway genes was tested for upregulation, downregulation and two-sided differential expression in comparison to the entire protein-coding genome. Enrichment of input genes for associated traits in GWAS catalogues was also tested in the FUMA framework.

### Individual profiling of *pharmagenic enrichment scores* in a genotyped schizophrenia cohort

We sought to generate PES for the schizophrenia candidate pathways in a cohort of diagnosed schizophrenia cases and non-neuropsychiatric controls sourced from the Australian Schizophrenia Research Bank (ASRB) ^17; 18^. Detailed descriptions of consent procedures along with inclusion and exclusion criteria for the ASRB have been extensively described elsewhere ^17^. The Illumina Infinitium Human 610K (610-Quad) BeadChip platform was used to genotype genomic DNA extracted from peripheral blood mononucleocytes as per standard manufacturer protocols. Variant and individual level quality control, along with imputation using the 1000 genomes phase 3 European reference panel, has been outlined in detail for this cohort previously ^18^. High quality autosomal sites with low missigness (< 2%) and an imputation score greater than 0.8 (R^2^ > 0.8) were retained for analysis in this study. After the removal of individuals in the post-genotyping quality control, 425 schizophrenia cases and 251 controls were analysed in this study; cases were 67% male, whilst males comprised 44% of the control cohort (Supplementary Table 5). The use of these data was approved by the University of Newcastle Human Ethics Research Committee (HREC).

PES is calculated from SNPs mapped to genes which form the candidate pharmacologically actionable geneset. This comprises the following model (1) which sums the statistical effect size of each variant in the geneset multiplied the allele count (dosage) for said variant– for individual *i*, let 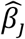 denote the statistical effect size from the GWAS for each variant *j* in the candidate gene-set, multiplied by the dosage (*G*) of *j* in *i*.

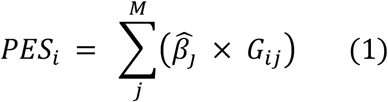

This was calculated for the individuals using the PRSice2 package ^19^. For each PES, the *P*_T_ used to derive the score was selected based on which *P*_T_ the corresponding geneset was derived from the GWAS, when a geneset was associated at multiple *P*_T_, the most significant was chosen. A genome-wide PRS (PRS_Total_) was also constructed for this cohort, with the *P*_T_ which explained the most variance between cases and controls selected using Nagelkerke’s R^2^. Association for each of the scores with cases was conducted using binomial logistic regression, adjusted for sex and the first three principal components using R 3.4.4. *P* values were derived using the Wald test with and without the total PRS score at the optimum threshold as a covariate in the model.

Each PES was ranked for individuals within the ASRB cohort, with three metrics used to define a person with an ‘elevated PES’ score: the top percentile, decile, and quartile of the study population. The number of pathways which pass these thresholds were totalled for each individual and the association between these totals and schizophrenia assessed using the same model as for univariate PRS as described above. To investigate the relationship between PRS_Total_ and PES, genome wide PRS_Total_ and the count of PES in the top decile per individual were clustered using finite Gaussian mixture modelling (GMM) with the mclust package version 5.4 ^20^. The optimal number of clusters was selected based on parametrisation of the covariance matrix utilising the Bayesian Information Criterion (BIC), with the highest BIC value used for selection of the number of clusters. Clusters were ellipsoidal, with the volume of the ellipsoid, shape of the density contours, and orientation of the corresponding ellipsoid also determined by the covariance matrix. We tested whether schizophrenia cases in each of the four GMM derived clusters were overrepresented for carriers of a top percentile PES using multinomial logistic regression with the nnet package (https://cran.r-project.org/web/packages/nnet/index.html). The largest cluster, *Cluster 2*, was used as the reference for the other clusters, with the model covaried for sex and prinicipal components as above. After dividing the regression coefficients by their standard error to derive *z, P* values were calculated using the Wald Test.

### Investigation of the effect of PES profiles on gene expression

We sought to investigate the relationship between PES profiles and the expression of genes which comprise their pathways in individuals with schizophrenia. A subset of schizophrenia cases in this cohort (N = 75) had mRNA expression data available from a previous study ^21^. These participants had a mean age of 42.21 (s.d. = 10.47), whilst the majority of the subcohort was male (N_Male_ = 45, N_Female_ = 30). RNA extracted from peripheral blood mononuclear cells was profiled using Illumina HT-12_V3 BeadChips and normalised as described in Gardiner *et al.* ^21^. Genes which comprise each PES pathway were extracted if they were available on the array with normalised expression values which survived quality control. The relationship between PES and the expression of each gene in that pathway as the outcome was assessed using a linear model covaried for age, sex, and PRS_Total_ for schizophrenia. These models were constructed in R version 3.4.4 using the *lm* function. Multiple testing correction was applied to each PES model individually to account for the number of genes tested in each pathway using Benjamini-Hochberg method via the *p.adjust* function in R.

## RESULTS

### Clinically actionable pathways enriched with common variant risk in schizophrenia

Schizophrenia is a typical complex trait disorder with a prevalence around 0.7% and heritability in the region of 80% ^22; 23^. A substantial proportion of this heritability was accounted for in the 2014 psychiatric genomics consortium (PGC) mega GWAS, which identified over 100 common variant loci at rigorous genome-wide significance level ^3^, making the disorder a suitable candidate to test the implementation of the PES framework. Using the complete summary statistics, we identified eight clinically actionable gene-sets (at the different *P*_T_) containing known drug targets (Table 1). The most significantly associated of these was the *HIF-2 pathway* (*P* = 3.12 × 10^-5^, *β* = 0.435, *SE* = 0.109, *P*_T_ < 0.005), which is comprised of genes in the hypoxia inducible factor 2 (HIF-2) alpha transcription factor network. *One carbon pool by folate* was the second most significant pathway with a putative drug interaction (*P* = 1.4 × 10^-4^, β= 0.433, *SE* = 0.119, *P*_T_ < 0.05). Two gene-sets were related to the function of the neurotransmitters GABA and Acetylcholine, whilst other signalling pathways represented were *NOS1* (Nitric Oxide Synthase I), *Hedgehog* signalling and the semaphorin related *CRMP (Collapsin Response Mediator) proteins in Sema3A signalling* pathway. In addition, the geneset *Regulation of Insulin Secretion* passed the threshold for inclusion.

**Table 1.** Pathways enriched with common variation associated with schizophrenia with putative clinical actionability. Pathways with putative clinical actionability by virtue of having targets for existing drugs with potential for repurposing. Enrichment *P* values refer gene-set association aggregated SNPs associated with schizophrenia in the PGC GWAS.

| Pathway | $P$ threshold ($P_T$) | $P$ |
| --- | --- | --- |
| NOS1 pathway | All SNPs | $6.3 \times 10^{-4}$ |
| Regulation of insulin secretion | $P < 0.5$ | $3.9 \times 10^{-4}$ |
| CRMPs in Sema3A signalling | $P < 0.5$ | $9.4 \times 10^{-4}$ |
| GABA synthesis, release, reuptake and degradation | $P < 0.5$ | $5.8 \times 10^{-4}$ |
| One carbon pool by folate | $P < 0.05$ | $1.4 \times 10^{-4}$ |
| Hedgehog signalling | $P < 0.05$ | $1.9 \times 10^{-4}$ |
| HIF-2 pathway | $P < 0.005$ | $3.1 \times 10^{-5}$ |
| Acetylcholine binding and downstream events | $P < 0.005$ | $3.8 \times 10^{-4}$ |

**Table 2.** The top enriched drug target for each *pharmagenic enrichment score* pathway with at least three interacting genes after multiple testing correction. Most significantly associated drug after multiple testing correction was selected, when corrected *P* values were equal, the drug with the highest geneset overlap was selected. Overlap refers to the number of genes targeted by the drug in the candidate pathway.

| Pathway | Top Drug* | Overlap<sup>#</sup> | ATC Code |
| --- | --- | --- | --- |
| NOS1 | Milnacipran | 5 | Other antidepressants |
| GABA | Temazepam | 14 | Benzodiazepine derivatives |
| Insulin | Clodronic acid | 3 | Bisphosphonates |
| HIF-2 | Ascorbic acid (Vitamin C) | 3 | Ascorbic acid (Vitamin C) |
| Acetylcholine | Nicotine | 11 | Drugs used in nicotine dependence |
| Folate | Tetrahydrofolic acid | 11 | Folic acid and derivatives |

The genes which constitute these eight pathways had upregulated expression in the brain relative to the rest of the protein coding genome, with the anterior cingulate cortex the most highly enriched region after multiple testing correction, *P*_Adj_ = 6.45 × 10^-13^. Conversely, they were downregulated (*P*_Adj_ < 0.05) in several peripheral tissues including the stomach and skin (Supplementary Fig. 2). These genes were also overrepresented in the GWAS catalogue for traits relevant to psychiatry including schizophrenia, post-traumatic stress disorder, nicotine dependence and cognitive performance (*P*_Adj_ < 0.05, Supplementary Table 1).

The eight gene-sets prioritised by our pipeline are indicative of a diverse range of drug classes. We sought to investigate a selection of candidate pharmacological agents which may be utilised for each PES input pathway. Firstly, we extracted the genes classified in the TCRD as T_clin_ from each of the gene-sets and matched them to their known drug-interactions using the drug gene interaction database (DGidb v3.02, Supplementary Table 2). The top FDA approved drug per pathway was selected based on the DGidb score of interaction confidence between a T_clin_ gene and drug. After annotation via the anatomical chemical (ATC) classification system two candidate drugs were anti-neoplastic and immunomodulating agents (ATC = L), two were classified as nervous system (ATC = N), whilst the remaining encompassed one of the following: blood and blood forming organs (ATC = B), musculoskeletal system (ATC = M), sensory organs (ATC = S) and alimentary tract and metabolism (ATC = A). Clinical trials for schizophrenia, either completed or in the recruiting phase, were registered for three of these compounds – glycine, varenicline and exenatide.

Drugs which target a statistically significant number of genes in each pathway were derived using over-representation analysis in WebGestalt ^14^. Of the eight gene-sets tested, six had a significant drug enrichment with a minimum overlap of three genes after multiple testing correction (Table 3, Supplementary Table 3). Nervous system drugs were the most common ATC category (level 1) across all the input pathways. Some interesting repurposing candidates with previous clinical trials in the disorder included the psychostimulant Atomoxiene ^24^, the α4β2 nicotinic acetylcholine receptor subtype partial agonist Varenicline ^25^, acetylcysteine (*N*-acetylcysteine) - a precursor to the antioxidant glutathione ^26–28^, ascorbic acid (Vitamin C) ^29; 30^, vitamin E^30^, and memantine ^31; 32^. Whilst the results of these trials were mixed, targeting such interventions to specific individuals based on genomic risk is yet to be investigated.

### Individual profiling of pharmagenic enrichment scores in a schizophrenia cohort

We profiled PES in a cohort of schizophrenia patients and screened healthy controls ^17^ and identified members of the cohort with relatively high PES in clinically actionable gene-sets. Firstly, we examined individuals in the top percentile of the ASRB cohort for each PES, to explore the phenotypic characteristics of an elevated risk score with high confidence. There were 55 individuals with a top percentile PES, as one schizophrenia case had elevated PES in both the *One Carbon Pool by Folate* and the *GABA synthesis, release, reuptake and degradation* pathways. From this subset, the majority were schizophrenia patients (N=38), however, there was no significant association between top percentile status and diagnosis (*z* = 0.975, *P* = 0.33). We investigated clinical characteristics obtained for ASRB participants to prioritise top percentile PES carriers who may benefit most from a personalised treatment regime. Three variables were selected as a proxy of a more clinically challenging presentation of the disorder: clozapine prescription (as a surrogate for treatment resistance), a global assessment of functioning (GAF) score < 50, and an adolescent onset of the disorder before the age of 18 ^33; 34^. Interestingly, of the 38 schizophrenia cases with an elevated PES, 71% of this subset meet at least one of these criteria (N =27): clozapine prescription (N = 9), GAF < 50 (N = 12), onset age < 18 (N = 9).

In addition, two less stringent partitions of elevated PES were implemented, specifically, a decile and quartile cut-off for PES in the entire cohort was used to triage patients at elevated risk of dysfunction in that pathway. The highest number of PES in the top decile or quartile respectively for an individual was six (Supplementary Fig. 3). An increasing number of PES in both the top quartile (OR = 1.1493 [95% CI: 1.016 – 1.303], *P* = 0.0287) and decile (OR = 1.207 [95% CI: 1.013 – 1.447], *P* = 0.0384) was associated with schizophrenia. However, this signal was not significant after adjustment for PRS_Total_. As visualised with kernel density estimation in figure 1b-c, there is evidence of skew towards high PRS_Total_ for those individuals with at least four top quartile or decile PES categories. Whilst the aim of this study was not to find association with cases for this cohort, two PES were nominally associated with schizophrenia in the ASRB - *Regulation of Insulin Secretion* (*z* = 2.262, *P* = 0.0237) and the *Acetylcholine Binding and Downstream Events* pathways (*z* = 2.167, *P* = 0.0303). However, significance was diminished when covaried for total schizophrenia PRS (PRS_Total_, *P* > 0.05).

**Fig 1.**
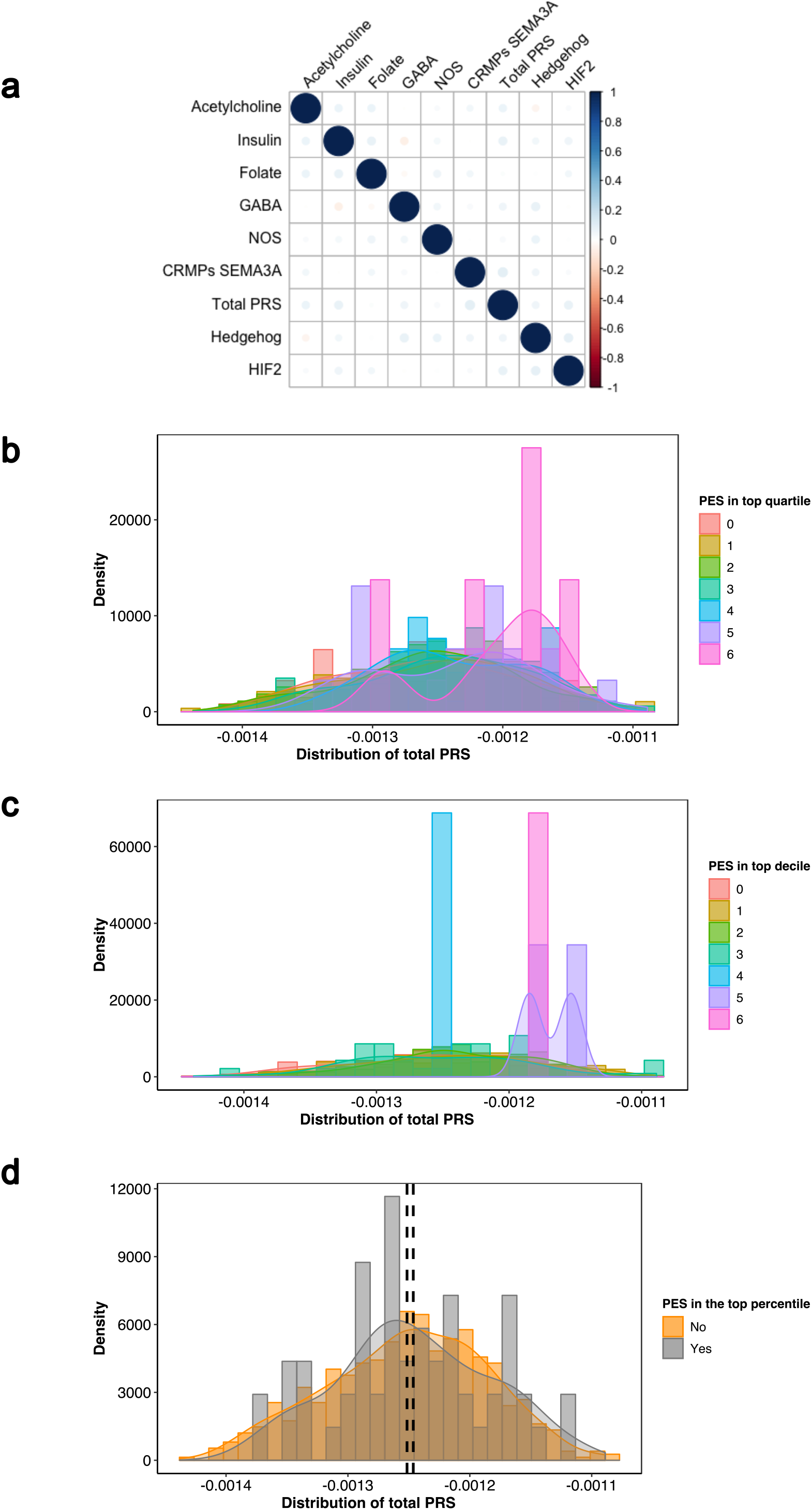
Relationship between genome wide schizophrenia PRS and *pharmagenic enrichment scores* in the ASRB cohort. (**a**) Pairwise univariate correlation between each of the PES and total PRS. Scale represents strength of relationship in the positive or negative direction. Kernel density estimation of distribution of total PRS amongst individuals with multiple PES in the top quartile (**b**) or decile (**c**). Scale refers to the number of PES over the threshold in an individual, that is, a score of six represents an individual with six PES categories in the top quartile or decile of the ASRB cohort. (**d**) Distribution of total PRS between ASRB participants with at least one PES in the top percentile of the cohort (grey) or without (orange). Black dashed line represents the mean PRS_Total_ for the cohort with a top percentile PES (right) and without (left).

### Relationship between pathway-based annotation and genome wide polygenic risk for schizophrenia

We sought to define the relationship between PRS_Total_ and PES in further detail. Pairwise correlation between each of the scores demonstrated no significant univariate relationship between any PES or with PRS_Total_ (Fig. 1a). In addition, top percentile PES individuals were not enriched with PRS_Total_ in comparison to the rest of the cohort: *z* = 0.819, *P* = 0.413 (Fig. 1d). This presented a clinically significant subset of schizophrenia cases with a less polygenic phenotype, that is, low PRS_Total_ relative to the schizophrenia cohort but high heritable risk in one or more pathways. Analysis of the bottom quartile of PRS_Total_ in the ASRB schizophrenia cohort revealed cases (N=10) with top percentile PES but depleted PRS_Total_. The pathways encompassed in these individuals were: *Acetylcholine* (N=3), *Hedgehog signalling* (N=2), *CRMPs in Sema3A* (N=2), *GABA* (N=1), *HIF-2* (N=1) and *Insulin secretion* (N=1). This information may be of great clinical value as these cases have less marked common variant burden genome-wide, but localised risk in a geneset. Furthermore, three of these patients were prescribed clozapine (surrogate for treatment resistance), a further three had low global functioning (GAF < 50), along with two adolescent onset cases – potentially highlighting a heightened need for precision intervention in these individuals.

To investigate the relationship between low polygenic load and elevated PES, genome wide PRS and the PES category count in the top decile per individual were clustered using finite Gaussian mixture modelling (GMM). The optimal number of clusters was selected based on parametrisation of the covariance matrix utilising the Bayesian Information Criterion (BIC), with the highest BIC value used for selection of the number of clusters (Fig. 2a). Four clusters were derived from the data (*BIC* = 13871.21, VEV: variable volume, equal shape, variable orientation) (Fig. 2b). Clusters were ellipsoidal, with the volume of the ellipsoid, shape of the density contours, and orientation of the corresponding ellipsoid also determined by the covariance matrix. The first two clusters were comprised of schizophrenia patients with at least one PES in the top decile of the ASRB cohort, with *Cluster 1* having low PRS_Total_ relative to *Cluster 2*. The third and fourth clusters had no elevated PES but *Cluster 3* represents patients with greater polygenic load, that is PRS_Total_, than the *Cluster 4*. The distribution of PRS_Total_ in *Cluster 1* reinforces the concept that a subset of schizophrenia patients with lower polygenic risk may have concentrated elevation in one or more specific biological systems. Analogous to its univariate relationship with PRS_Total_ individuals with extreme PES in the top percentile of the ARSB cohort were not enriched in any of the GMM clusters relative to the largest cluster, *Cluster 2* (*Cluster 2* vs *Cluster 1*: *P* = 0.668; *Cluster 2* vs *Cluster 3*: *P* = 0.492; *Cluster 2* vs *Cluster 4*: *P* = 0.977).

**Fig 2.**
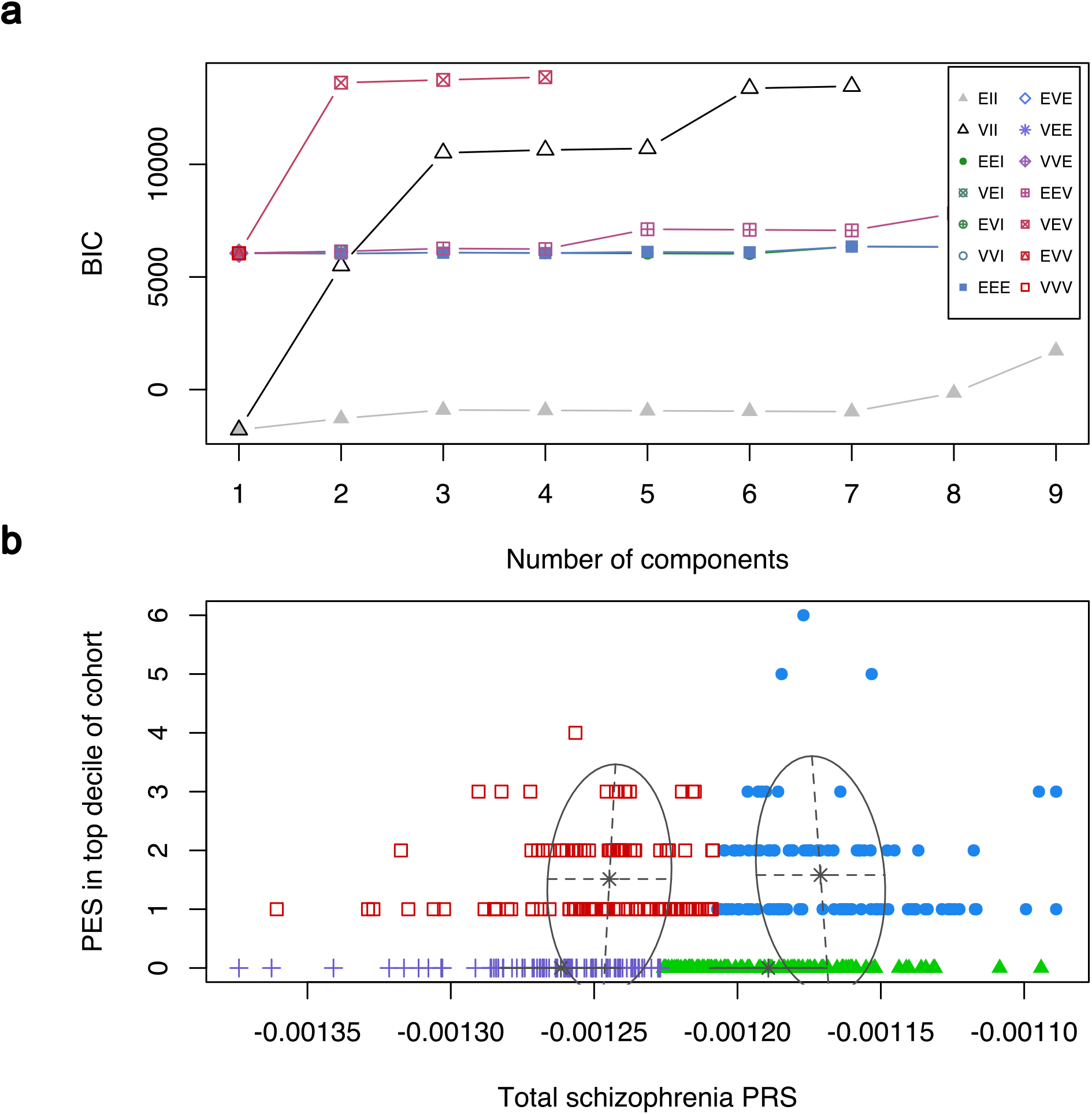
Clustering of genome wide PRS and the individual count of elevated *pharmagenic enrichment score* using finite Gaussian mixture modelling. (**a**) Selection of the number of clusters using the Bayesian information criterion (BIC). The scale represents the fourteen different Gaussian models (see Supplementary Table 5 for definitions) tested for parametrisation of the within-group covariance matrix. (**b**) Derived clusters of genome wide schizophrenia PRS (Total PRS) and the number of PES in the top decile of the ASRB cohort per individual. Red boxes = *Cluster 1*, blue circles = *Cluster 2*, green triangles = *Cluster* 3, purple crosses = *Cluster 4*.

### PES profiles impact the expression of genes within the candidate pathways

Each PES profile was tested for association with the peripheral blood expression of genes within their respective pathways for a subset of schizophrenia cases with expression data available (Fig. 3, Supplementary Table 6). After covariation for sex, age, and PRS_Total_, the *NOS1* PES was associated with downregulated expression of the calcineurin subunit gene *PPP3CC* (*t* = −3.08, *P* = 2.9 × 10^-3^, *q* = 0.05). This was followed by the *regulation of insulin secretion* PES, which was associated with decreased expression of the syntaxin gene *STX1A* (*t* = −3.5, *P* = 8.1 × 10^-4^, *q* = 0.055); and the *One carbon pool by folate gene* PES, which was associated with downregulation of serine hydroxymethyltransferase 2 (*SHMT2*; *t* = −2.94, *P* = 4.5 × 10^-3^, *q* = 0.072). Excluding these three genes (*q* < 0.1), there were eleven others with a nominally significant (Raw *P* < 0.05) relationship with a PES, with all PES profiles except *CRMPs in Sema3A signalling* having at least one such nominally significant model.

**Figure 3.**
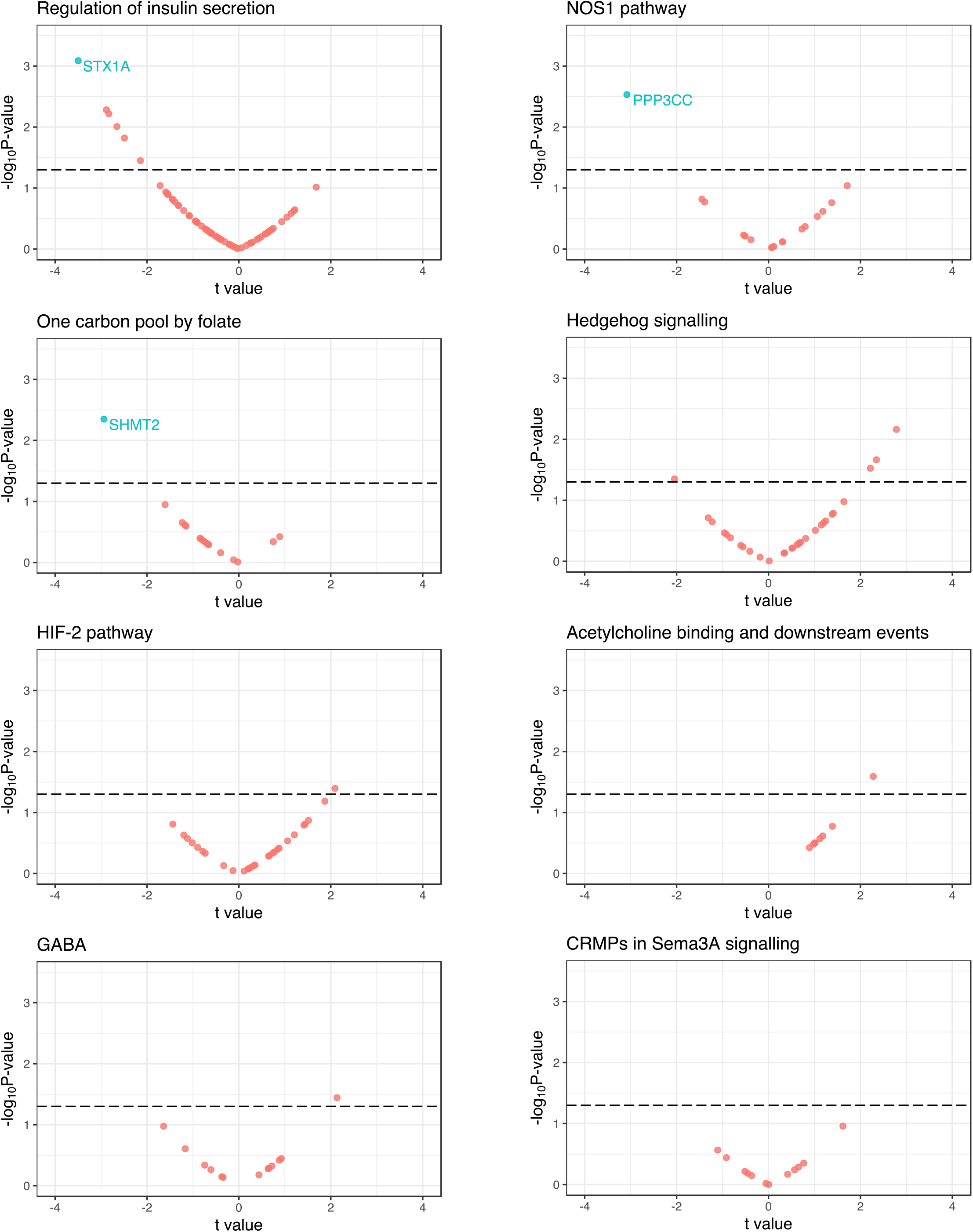
The relationship between PES profiles and the expression of pathway genes in peripheral blood mononuclear cells. Plot for each PES of the results of a model which investigated the association of PES with the expression of genes which compromise the PES pathway. The *t* values on the x-axes were derived from the regression model for each PES-gene pair 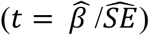, with the −log_10_ *P* value of association on the y-axes. The dotted line is indicative of uncorrected *P* < 0.05, genes highlighted blue were significant after correcting for the number of genes in the set with a liberal false discovery rate cut-off (*q* < 0.1).

## DISCUSSION

The drug development pipeline continues to be prohibitively expensive and time consuming in the translation of novel compounds for clinical practice ^35; 36^. Repositioning of previously approved drugs for other human health conditions can be a more readily achievable action, particularly for rare disorders where a causal factor can be identified. However, in complex disorders, such as schizophrenia, this approach is hindered by the complexity of the pathophysiology and heterogeneity of genomic risk, along with inter-individual variability in illness onset and clinical course ^37; 38^. Annotation of the individually-relevant (personalised) genetic components associated with complex syndromes, for the purpose of delineating clinically meaningful biological systems, will both better target existing treatments and reveal new opportunities for drug repurposing (Fig. 4a). In this study, we developed a novel method for capturing common variant risk in biological networks with known drug interactions – *pharmagenic enrichment scores* (PES) – to facilitate precision treatment design relevant to individuals with a particular set of risk variants. A distinct advantage of our PES approach is that it can capture latent enrichment of polygenic signal in pathways relevant to pharmaceutical actions, among individuals whose overall trait PRS is low relative to others with a shared phenotype. Even in cases where polygenic burden is high, genome-wide PRS (as a biologically unannotated instrument) does not necessarily provide insight into pathways suitable for pharmacological intervention in individuals. Our approach for selection of putative drug targets exemplified in schizophrenia GWAS has revealed potential targets for drug repurposing with substantial clinical utility.

**Fig. 4.**
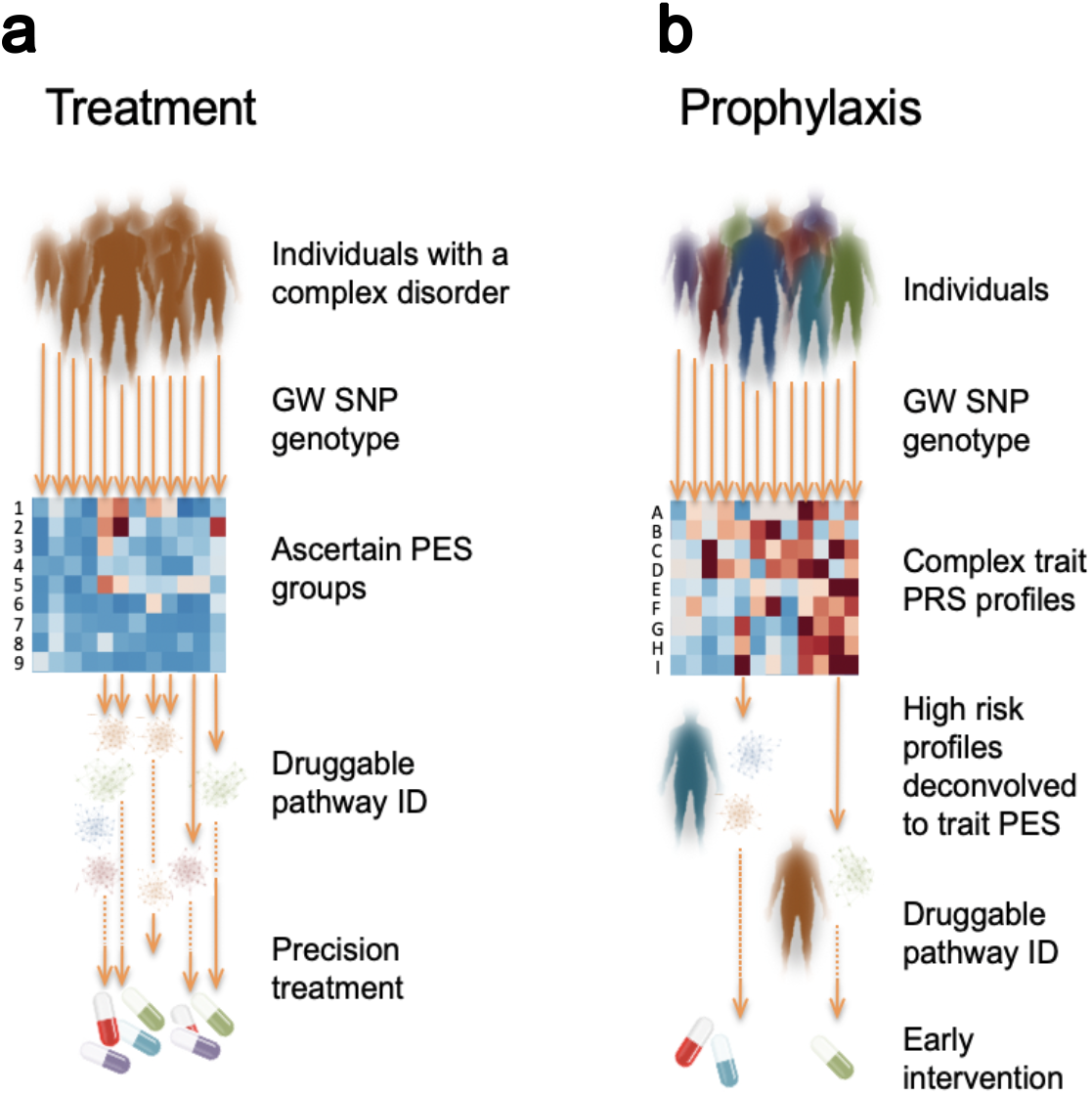
Implementation of *pharmagenic enrichment score* (PES) in precision treatment and prophylaxis of complex disorders. (**a**) Using the PES framework individuals with a complex disorder provide DNA for common variant SNP genotyping, which is used to ascertain individuals with a high PES. The PES groups (heatmap rows 1-9, with enrichment denoted by darker colour) intrinsically identify precision treatment options tailored to the individual’s biological enrichment (of pathways with known drug targets) for polygenic risk in that pathway. (**b**) A more advanced implementation of PES could be achieved for prophylactic intervention for individuals in the population at very high polygenic risk for a variety of complex traits with clinical actionability (heatmap rows A-I, with high PRS denoted by darker colour). Conservative prophylactic measures in this context may account for environmental risk exposure and focus on lifestyle interventions, such as diet and exercise, rather than pharmaceutical treatments that may not be justified without symptom presentation because of their side effect and/or cost.

Aggregation of common variation from schizophrenia GWAS into biological pathways with known drug interactions revealed a diverse array of systems relevant to eight distinct PES categories. These candidate pathways displayed common variant enrichment at a range of *P*_T_, indicative of the degree of polygenicity, ranging from using all SNPs as input, to a significance threshold below 0.005 (*P*_T_ < 0.005). While two of these pathways included GABAergic and cholinergic neurotransmission, both of which are intuitive candidates that have been extensively implicated in schizophrenia with associated drugs already in common practice for neuropsychiatric disorders ^39; 40^, many others were more surprising. The most significantly associated gene-set pertained to the HIF-2 transcription factor network, an important mediator in response to decreases in available cellular oxygen. This has clear significance for biological mechanisms involved in psychiatric disorders, for example in dopaminergic signalling ^41^. Enrichment of ascorbic acid (vitamin C) targets in this pathway is notable from a therapeutic perspective because of its antioxidant capabilities, along with preliminary evidence for its efficacy as an adjuvant in the treatment of the disorder ^29; 30^. The interaction between HIF-2 signalling and NOS1 signalling, another candidate pathway with pharmagenic enrichment in schizophrenia, is supported by previous evidence of redox dysfunction in the disorder ^42; 43^. The activity of glutamate receptors in the NOS1 system suggests that psycholeptics and psychoanaleptics are likely to modulate this pathway. We also observed common variant enrichment in two developmental pathways that can be pharmacologically modulated: CRMPs in semaphorin 3a signalling and Hedgehog signalling.

The former is able to interact with the tyrosine kinase inhibitor Dasatinib, which is postulated to have neuroprotective properties ^44^. Enrichment in these actionable pathways is consistent with longstanding hypotheses of deficits in neurodevelopment contributing to the aetiology of schizophrenia ^45^. There is also evidence for aetiological overlap between schizophrenia and diabetes beyond what is attributable to metabolic effects of antipsychotic treatment, which supports our identification of an insulin related pathway as a candidate PES ^46–48^.

The breadth of drugs which target these pathways used to construct PES suggests that individual level treatment formulation can become highly specific depending on which systems genomic risk is localised. This would include the stratification of individuals for precision treatment with compounds previously tested on undifferentiated schizophrenia cohorts, including, N-acetylcysteine, vitamin C, Atomoxiene, and Varenicline which were identified using PES in this study ^24–26; 29^. Repurposing drugs for individuals informed by their genetic liability may assist in the reduction of response heterogeneity, which hinders the implementation of novel treatments in very complex phenotypes like schizophrenia. We suggest that the individuals with PES in the top percentile of any pathway, particularly those with low genome wide PRS, present as the most tractable candidates for this approach; whereas the clinical significance of particular sets of common variant burden would be missed by an unannotated genome wide association indexed by total PRS alone.

In order to better understand how PES profiles could be leveraged for treatment, the effect of sequence variation which comprises the PES needs to be investigated. We outlined an example of this approach in this study by testing the effect of PES on the expression of genes which comprise each pathway in schizophrenia. Several associations between PES and mRNA expression were observed after correcting for the number of genes within the tested set. These effects may arise due to direct *cis*-acting loci within the PES and/or the downstream biologic effects of variation which effect genes with interrelated functionality. This was exemplified by the *Regulation of insulin secretion* PES, which was negatively correlated with *STX1A* expression, a syntaxin postulated to play a role in insulin homeostasis. Interestingly, *STX1A* has been shown to be positively correlated with glucose stimulated insulin secretion, suggesting downregulation conferred by the PES may have an important effect through this biological pathway^49; 50^. Similarly, downregulation of the calcineurin subunit gene *PPP3CC* was associated with the *NOS1 pathway* PES. Previous research suggests there is a bidirectional relationship between nitric oxide signalling and calcineurin, where the calcineurin subunit is both regulated by redox products and able to induce nitric oxide synthesis ^51; 52^. Several other genes had suggestive association with PES profiles which may be established with a larger cohort. We suspect that this approach to validation would be particularly informative for schizophrenia in genotyped expression cohorts from brain tissue.

While we expect that in most circumstances the aggregate of variation constituting high PES represent pathology of the target pathways, a current limitation of this methodology is that it does not integrate the direction of effect. While this may be possible as more functional annotations become available, this would present an immensely complex paradigm to predict *in silico* due to the vast array of factors which influence the penetrance of genomic risk. We believe that an analysis of the effect of PES profiles on gene expression in larger cohorts will be an important future direction of this work. The impact of candidate molecules could also be modelled in patient derived cell lines to, firstly, establish the extent of dysregulation conferred by elevated PES and, secondly, investigate the interaction with the compounds implicated. The current analysis also used very stringent criteria for drug pathway gene inclusion and there are likely to be many more genes and pathways that may be implicated in future ontologies and other investigator curated gene-sets. In a recent example of this we observed an enrichment of retinoid gene variation in schizophrenia ^18^.

New GWAS summary statistics are emerging daily on ever larger samples and these too will further enrich the substrate for PES determination and increase its clinical utility. Despite the aforementioned challenges, we believe that this methodology provides a useful framework to better utilise the breadth of available GWAS data for personalised treatment formulations. Particularly, as there remains a largely unmet need to translate polygenic risk for complex disorders into tractable treatment outcomes for affected individuals. Whilst we have demonstrated here the potential utility in schizophrenia, there is clearly scope to adapt this analytical approach to other complex disorders with summary statistics from well-powered GWAS. This methodology may also be applicable to prophylactic intervention for individuals at high risk for a complex phenotype (Fig. 4b). This could be implemented conservatively with lifestyle or dietary measures implicated by clusters of enrichment captured within the PES framework. For example, in schizophrenia we identified multiple actionable pathways quantified by PES that are modulated by vitamins, which represent a relatively uncomplicated intervention for individuals at high genetic risk for this disorder.

## SUPPLEMENTAL DATA

**Supplementary Fig 1.** Methodology for identifying pharmacologically-relevant pathways enriched with GWAS risk variants.

**Supplementary Fig 2.** Tissue specific expression of genes contained within candidate PES pathways derived from schizophrenia GWAS

**Supplementary Fig. 3.** Distribution of schizophrenia and healthy control patients with multiple elevated *pharmagenic enrichment scores*.

**Supplementary Table 1.** Overrepresentation of genes within candidate PES pathways in the GWAS catalogue traits with relevance to psychiatry after multiple testing correction.

**Supplementary Table 2.** Highest confidence drug interaction between of a member of each pathway enriched with common polygenic risk for schizophrenia.

**Supplementary Table 3.** Enriched drug targets for each *pharmagenic enrichment score* with at least three interacting genes after multiple testing correction (FDR < 0.05).

**Supplementary Table 4.** Geometric characteristics of the Gaussian models used for parameterisations of the within-group covariance matrix.

**Supplementary Table 5.** Characteristics of the ASRB cohort analysed using the PES methodology.

**Supplementary Table 6.** Effect of PES on gene expression for genes which comprise each of the candidate pathways.

## DECLERATIONS OF INTERESTS

The authors declare no conflicting financial interests

## DATA AVAILABILITY

Schizophrenia GWAS data is available from the Psychiatric Genomics Consortium (https://www.med.unc.edu/pgc/results-and-downloads). SNP array data used in this study is available upon application to the Australian Schizophrenia Research Bank, URL: https://www.neura.edu.au/discovery-portal/asrb/. Command line arguments for the bioinformatics tools in this study are available upon reasonable request to the authors.

