## Supplemental Material for "Pharmacological enrichment of polygenic risk for precision medicine in complex disorders"

**Supplementary Materials**

**Supplementary Fig 1.** Methodology for identifying pharmacologically-relevant pathways enriched with GWAS risk variants.

**Supplementary Fig 2.** Tissue specific expression of genes contained within candidate PES pathways derived from schizophrenia GWAS

**Supplementary Fig. 3.** Distribution of schizophrenia and healthy control patients with multiple elevated *pharmagenic enrichment scores*.

**Supplementary Table 1.** Overrepresentation of genes within candidate PES pathways in the GWAS catalogue traits with relevance to psychiatry after multiple testing correction.

**Supplementary Table 2.** Highest confidence drug interaction between of a member of each pathway enriched with common polygenic risk for schizophrenia.

**Supplementary Table 3.** Enriched drug targets for each *pharmagenic enrichment score* with at least three interacting genes after multiple testing correction (FDR < 0.05).

**Supplementary Table 4**. Geometric characteristics of the Gaussian models used for parameterisations of the within-group covariance matrix.

**Supplementary Table 5.** Characteristics of the ASRB cohort analysed using the PES methodology.

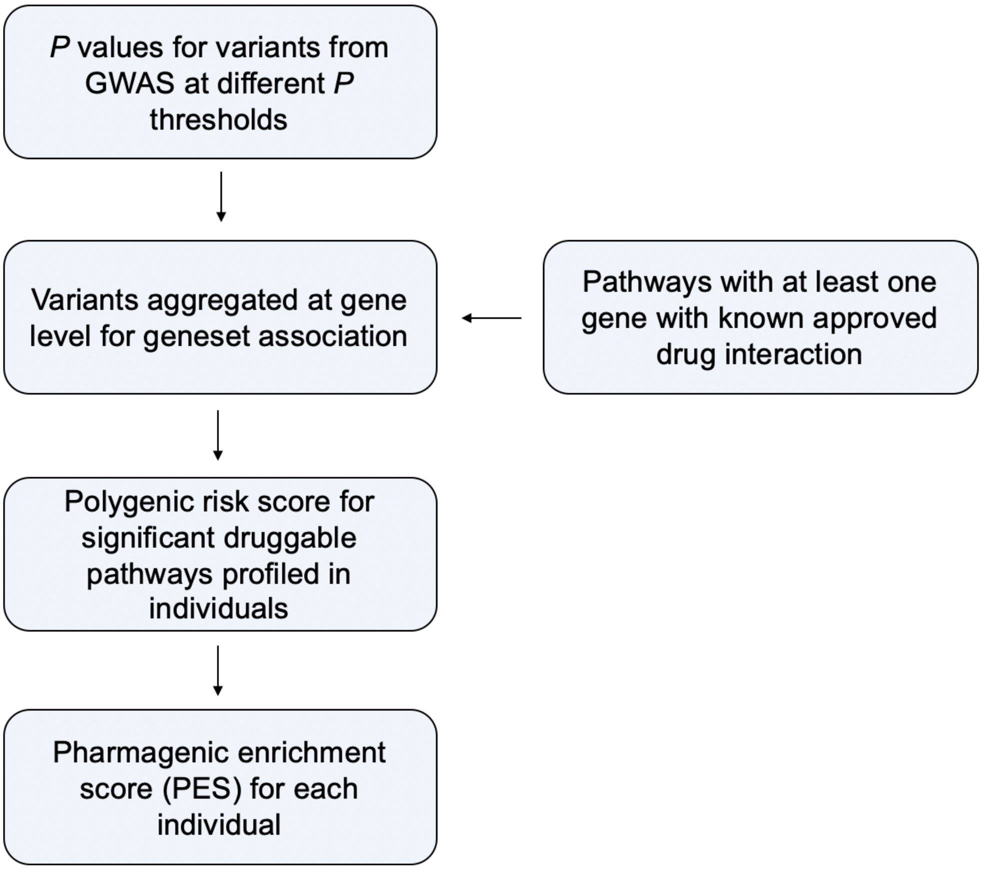


**Supplementary Fig 1. Methodology for identifying pharmacologically-relevant pathways enriched with GWAS risk variants.** The combined effect of variants (*P* values) from a genome wide association study (GWAS) is tested at level of genes. Different *P* value thresholds ($P_T$) are used to filter variants for input to capture biological signals only present at varying levels of polygenicity. Using a regression approach implemented by the MAGMA algorithim, geneset association is undertaken at each $P_T$ and pathways are then filtered based on likelihood of interaction with an approved drug. Polygenic risk score (PRS) is constructed for each significant pathway uncovered via the pipeline to formulate a *pharmagenic enrichment score (PES)*. This PES can then be profiled in individuals to reveal participants with elevated PES, which in turn may be relevant to treatment formulation.

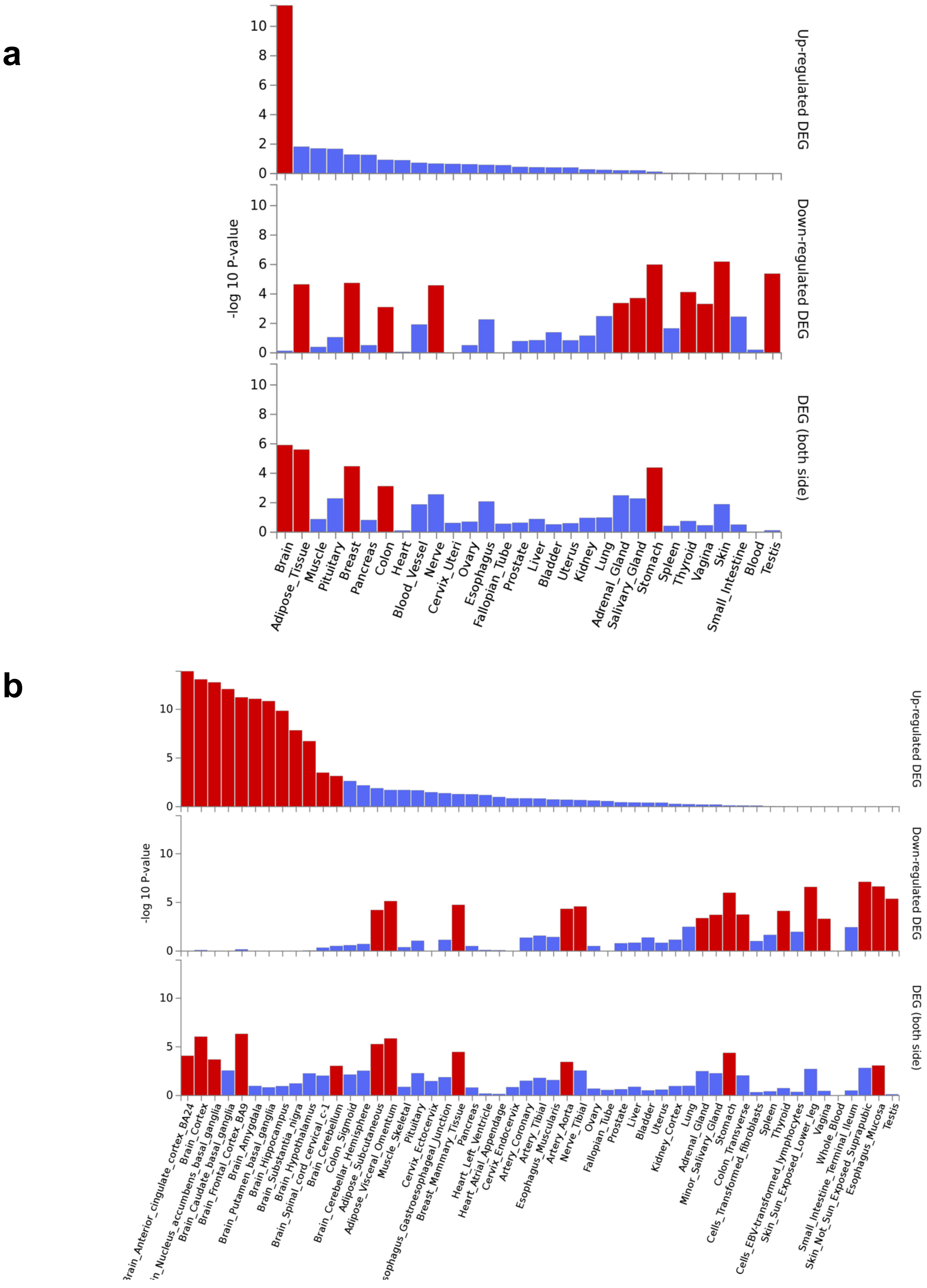
a
b
-log 10 P-value
Up-regulated DEG
Down-regulated DEG
DEG (both side)
0
2
4
6
8
10
5
Brain
Adipose_Tissue
Muscle
Pituitary
Breast
Pancreas
Colon
Heart
Blood_Vessel
Nerve
Cervix_Uteri
Ovary
Esophagus
Fallopian_Tube
Prostate
Liver
Bladder
Uterus
Kidney
Lung
Adrenal_Gland
Salivary_Gland
Stomach
Spleen
Thyroid
Vagina
Skin
Small_Intestine
Blood
Testis
Brain_Cortex
Brain_Caudate_basal_ganglia
Brain_Frontal_Cortex_BA9
Brain_Amygdala
Brain_Putamen_basal_ganglia
Brain_Hippocampus
Brain_Substantia_nigra
Brain_Hypothalamus
Brain_Spinal_cord_cervical_c-1
Brain_Cerebellum
Colon_Sigmoid
Brain_Cerebellar_Hemisphere
Adipose_Subcutaneous
Adipose_Visceral_Omentum
Muscle_Skeletal
Pituitary
Cervix_Ectocervix
Breast_Mammary_Tissue
Pancreas
Heart_Left_Ventricle
Heart_Atrial_Appendage
Cervix_Endocervix
Artery_Coronary
Artery_Tibial
Esophagus_Muscularis
Artery_Aorta
Nerve_Tibial
Ovary
Fallopian_Tube
Prostate
Liver
Bladder
Uterus
Kidney_Cortex
Lung
Adrenal_Gland
Minor_Salivary_Gland
Stomach
Colon_Transverse
Cells_Transformed_fibroblasts
Spleen
Thyroid
Cells_EBV-transformed_lymphocytes
Skin_Sun_Exposed_Lower_leg
Vagina
Whole_Blood
Small_Intestine_Terminal_Ileum
Skin_Not_Sun_Exposed_Suprapubic
Esophagus_Mucosa
Testis

**Supplementary Fig 2. Tissue specific expression of genes contained within candidate PES pathways derived from schizophrenia GWAS.** Expression per tissue for genes which comprise these pathways was compared to the rest of the protein coding genome, with the -$\log_{10}$(*P*-value) reported for each test after the application of multiple testing correction (red bars indicating tissues which survive correction). Tissue specific expression was performed to assess up-regulation, downregulation and a two-sided test of differential expression. (**a**) GTEx v7 30 tissue types. (**b**) GTEx v7 53 tissue types.

**a**

**b**

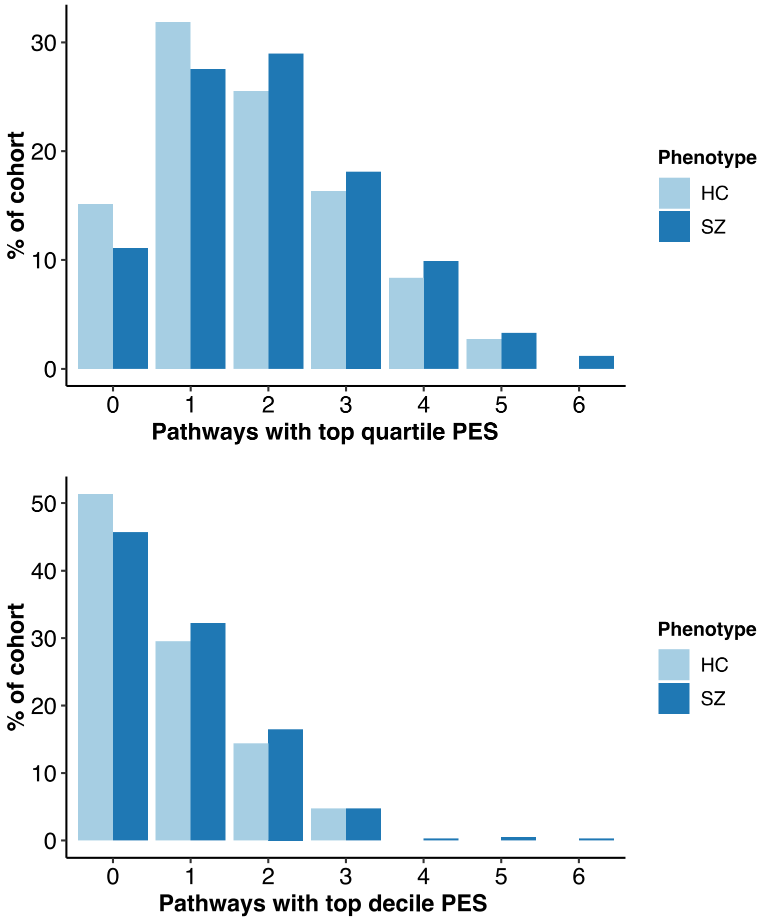


**Supplementary Fig 3. Distribution of schizophrenia and healthy control patients with multiple elevated *pharmagenic enrichment scores*.** PES in each pathway in the top quartile (**a**) or decile (**b**) of the ASRB cohort are classified as high and the number of scores over this threshold counted in each individual, represented here as a percentage of each phenotype cohort (that is, SZ or HC). The highest number of top quartile or decile PES scores in an individual is six (only schizophrenia patients). SZ = schizophrenia, HC = healthy controls.

| **Phenotype** | **Overlap** | **Adjusted *P*-value** |
|---|---|---|
| Nicotine dependence | 7 | $4.28 \times 10^{-8}$ |
| Smoking behaviour | 5 | $8.95 \times 10^{-5}$ |
| Schizophrenia | 18 | 0.0023 |
| Excessive daytime sleepiness | 2 | 0.017 |
| Post-traumatic stress disorder | 2 | 0.021 |
| Hippocampal volume | 2 | 0.024 |
| PGC cross disorder | 3 | 0.033 |
| Cognitive performance | 4 | 0.034 |
| White matter hyperintensity burden | 2 | 0.035 |
| Night sleep phenotypes | 10 | 0.036 |
| Social communication problems | 2 | 0.044 |
| Cerebral amyloid deposition in APOEε4 non-carriers (PET imaging) | 2 | 0.044 |

**Supplementary Table 1.** Overrepresentation of genes within candidate PES pathways in the GWAS catalogue traits with relevance to psychiatry after multiple testing correction. Overlap column pertains to genes common to the PES geneset and the trait geneset

| Pathway | Drug* | ATC Code | ATC Code Level 4 |
|---|---|---|---|
| NOS1 | Glycine | B05CX03 | Other irrigating solutions |
| GABA | Baclofen | M03BX01 | Other centrally acting agents# |
| CRMPs Sema3A | Dasatinib | L01XE06 | Protein kinase inhibitors |
| HIF-2 | Sunitinib | L01XE04 | Protein kinase inhibitors |
| Acetylcholine | Varenicline | N07BA03 | Drugs used in nicotine dependence |
| Hedgehog | Tacrine | N06DA01 | Anticholinesterases |
| Folate | Trifluridine | S01AD02 | Antivirals |
| Insulin | Exenatide | A10BJ01 | Glucagon-like-peptide-1 (GLP-1) analogue |

**Supplementary Table 2. Highest confidence drug interaction between of a member of each pathway enriched with common polygenic risk for schizophrenia**. Drugs selected by highest confidence interaction score with a gene (classified as $T_{Clin}$) in each of the pathways by DGidb v3.02

| PES PATHWAY | DRUGBANK ID | DRUG NAME |
|---|---|---|
| NOS1 | DB06741 | Gavestinel |
| NOS1 | DB04896 | Milnacipran |
| NOS1 | DB00289 | Atomoxetine |
| NOS1 | DB06738 | Ketobemidone |
| NOS1 | DB00996 | Gabapentin |
| NOS1 | DB01174 | Phenobarbital |
| NOS1 | DB01520 | Tenocyclidine |
| NOS1 | DB00454 | Pethidine |
| NOS1 | DB00418 | Secobarbital |
| NOS1 | DB02868 | 3"-(Beta-Chloroethyl)-2",4"-Dioxo-3, 5"-Spiro-Oxazolidino-4-Deacetoxy-Vinblastine |
| NOS1 | DB06151 | Acetylcysteine |
| NOS1 | DB00659 | Acamprosate |
| NOS1 | DB00312 | Pentobarbital |
| NOS1 | DB01429 | Aprindine |
| NOS1 | DB03977 | N-Trimethyllysine |
| NOS1 | DB04825 | Prenylamine |
| NOS1 | DB08039 | (3Z)-N,N-DIMETHYL-2-OXO-3-(4,5,6,7-TETRAHYDRO-1H-INDOL-2-YLMETHYLIDENE)-2,3-DIHYDRO-1H-INDOLE-5-SULFONAMIDE |
| NOS1 | DB01708 | Dehydroepiandrosterone |
| NOS1 | DB00527 | Cinchocaine |
| NOS1 | DB00623 | Fluphenazine |
| NOS1 | DB00850 | Perphenazine |
| NOS1 | DB04513 | N-(6-Aminohexyl)-5-Chloro-1-Naphthalenesulfonamide |
| NOS1 | DB01043 | Memantine |
| NOS1 | DB01100 | Pimozide |
| NOS1 | DB03900 | 2-Methyl-2-Propanol |
| NOS1 | DB04841 | Flunarizine |
| NOS1 | DB08231 | Myristic acid |
| NOS1 | DB00831 | Trifluoperazine |
| NOS1 | DB00836 | Loperamide |
| NOS1 | DB00925 | Phenoxybenzamine |
| NOS1 | DB01115 | Nifedipine |
| NOS1 | DB01065 | Melatonin |
| NOS1 | DB01244 | Bepridil |
| NOS1 | DB01373 | Calcium |
| NOS1 | DB01069 | Promethazine |
| NOS1 | DB00142 | L-Glutamic Acid |
| NOS1 | DB01023 | Felodipine |
| NOS1 | DB00949 | Felbamate |

| | | |
|---|---|---|
| NOS1 | DB02527 | Cyclic Adenosine Monophosphate |
| NOS1 | DB00622 | Nicardipine |
| NOS1 | DB00753 | Isoflurane |
| NOS1 | DB00477 | Chlorpromazine |
| NOS1 | DB01173 | Orphenadrine |
| NOS1 | DB00163 | Vitamin E |
| GABA | DB00186 | Lorazepam |
| GABA | DB00189 | Ethchlorvynol |
| GABA | DB00228 | Enflurane |
| GABA | DB00231 | Temazepam |
| GABA | DB00237 | Butabarbital |
| GABA | DB00241 | Butalbital |
| GABA | DB00273 | Topiramate |
| GABA | DB00292 | Etomidate |
| GABA | DB00306 | Talbutal |
| GABA | DB00312 | Pentobarbital |
| GABA | DB00349 | Clobazam |
| GABA | DB00371 | Meprobamate |
| GABA | DB00402 | Eszopiclone |
| GABA | DB00404 | Alprazolam |
| GABA | DB00463 | Metharbital |
| GABA | DB00475 | Chlordiazepoxide |
| GABA | DB00546 | Adinazolam |
| GABA | DB00628 | Clorazepate |
| GABA | DB00659 | Acamprosate |
| GABA | DB00683 | Midazolam |
| GABA | DB00690 | Flurazepam |
| GABA | DB00753 | Isoflurane |
| GABA | DB00794 | Primidone |
| GABA | DB00801 | Halazepam |
| GABA | DB00818 | Propofol |
| GABA | DB00829 | Diazepam |
| GABA | DB00842 | Oxazepam |
| GABA | DB00897 | Triazolam |
| GABA | DB01028 | Methoxyflurane |
| GABA | DB01049 | Ergoloid |
| GABA | DB01068 | Clonazepam |
| GABA | DB01107 | Methyprylon |
| GABA | DB01159 | Halothane |
| GABA | DB01189 | Desflurane |
| GABA | DB01205 | Flumazenil |
| GABA | DB01215 | Estazolam |
| GABA | DB01236 | Sevoflurane |
| GABA | DB01437 | Glutethimide |
| GABA | DB01558 | Bromazepam |

| | | |
|---|---|---|
| GABA | DB01559 | Clotiazepam |
| GABA | DB01567 | Fludiazepam |
| GABA | DB01588 | Prazepam |
| GABA | DB01589 | Quazepam |
| GABA | DB01594 | Cinolazepam |
| GABA | DB01595 | Nitrazepam |
| GABA | DB01708 | Dehydroepiandrosterone |
| GABA | DB11582 | Thiocolchicoside |
| GABA | DB00543 | Amoxapine |
| GABA | DB00334 | Olanzapine |
| GABA | DB05087 | Ganaxolone |
| GABA | DB00898 | Ethanol |
| GABA | DB00849 | Methylphenobarbital |
| GABA | DB01351 | Amobarbital |
| GABA | DB01352 | Aprobarbital |
| GABA | DB01353 | Butethal |
| GABA | DB01354 | Heptabarbital |
| GABA | DB01355 | Hexobarbital |
| GABA | DB01483 | Barbital |
| GABA | DB01496 | Barbituric |
| GABA | DB00599 | Thiopental |
| GABA | DB01544 | Flunitrazepam |
| GABA | DB00418 | Secobarbital |
| GABA | DB01198 | Zopiclone |
| GABA | DB00425 | Zolpidem |
| GABA | DB01381 | Ginkgo |
| GABA | DB01587 | Ketazolam |
| GABA | DB00466 | Picrotoxin |
| GABA | DB01346 | Quinidine |
| Insulin | DB00720 | Clodronate |
| Insulin | DB00661 | Verapamil |
| HIF-2 | DB00126 | Vitamin C |
| Acetylcholine | DB00184 | Nicotine |
| Acetylcholine | DB00674 | Galantamine |
| Acetylcholine | DB00898 | Ethanol |
| Acetylcholine | DB05740 | RPI-78M |
| Acetylcholine | DB01273 | Varenicline |
| Acetylcholine | DB09028 | Cytisine |
| Acetylcholine | DB00514 | Dextromethorphan |
| Acetylcholine | DB00849 | Methylphenobarbital |
| Acetylcholine | DB01351 | Amobarbital |
| Acetylcholine | DB01352 | Aprobarbital |
| Acetylcholine | DB01353 | Butethal |
| Acetylcholine | DB01354 | Heptabarbital |
| Acetylcholine | DB01355 | Hexobarbital |
| Acetylcholine | DB01483 | Barbital |

| | | |
|---|---|---|
| Acetylcholine | DB01496 | Barbituric acid |
| Acetylcholine | DB00599 | Thiopental |
| Acetylcholine | DB01174 | Phenobarbital |
| Acetylcholine | DB01090 | Pentolinium |
| Acetylcholine | DB01227 | Levomethadyl |
| Acetylcholine | DB00418 | Secobarbital |
| Acetylcholine | DB00237 | Butabarbital |
| Acetylcholine | DB00241 | Butalbital |
| Acetylcholine | DB00306 | Talbutal |
| Acetylcholine | DB00463 | Metharbital |
| Acetylcholine | DB00794 | Primidone |
| Acetylcholine | DB00312 | Pentobarbital |
| Folate | DB00116 | Tetrahydrofolic acid |
| Folate | DB00642 | Pemetrexed |

**Supplementary Table 3.** Enriched targets for each pharmagenic enrichment score pathway with at least three interacting (overlapping) genes after multiple testing correction (FDR adjusted $P < 0.05$).

| Model abbreviation* | Distribution | Volume | Shape | Orientation |
|---|---|---|---|---|
| EEI | Spherical | Equal | Equal | - |
| VII | Spherical | Variable | Equal | - |
| EEI | Diagonal | Equal | Equal | Coordinate axes |
| VEI | Diagonal | Variable | Equal | Coordinate axes |
| EVI | Diagonal | Equal | Variable | Coordinate axes |
| VVI | Diagonal | Variable | Variable | Coordinate axes |
| EEE | Ellipsoidal | Equal | Equal | Equal |
| EVE | Ellipsoidal | Equal | Variable | Equal |
| VEE | Ellipsoidal | Variable | Equal | Equal |
| VVE | Ellipsoidal | Variable | Variable | Equal |
| EEV | Ellipsoidal | Equal | Equal | Variable |
| VEV | Ellipsoidal | Variable | Equal | Variable |
| EVV | Ellipsoidal | Equal | Variable | Variable |
| VVV | Ellipsoidal | Variable | Variable | Variable |

**Supplementary Table 4.** Geometric characteristics of the Gaussian models used for parameterisations of the within-group covariance matrix. Gaussian models described as implemented in the mclust package.

| | Control | Case |
|---|---|---|
| Total | 251 | 425 |
| Males | 110 | 283 |
| Females | 141 | 142 |
| Mean Age (s.d.) | 39.50 (13.40) | 39.88 (10.92) |
| Mean Onset Age (s.d.) | N/A | 23.79 (6.89) |
| Mean GAF score (s.d.) | 84.13(8.81) | 53.56 (13.43) |

**Supplementary Table 5.** Characteristics of the ASRB cohort to which the PES pipeline was applied to – cases refer to subjects diagnosed with schizophrenia. GAF refers to the global assessment of functioning scale, with a lower score indicating greater symptom severity for the disorder.
